# Identification of TMEM106B as proviral host factor for SARS-CoV-2

**DOI:** 10.1101/2020.09.28.316281

**Authors:** Jim Baggen, Leentje Persoons, Sander Jansen, Els Vanstreels, Maarten Jacquemyn, Dirk Jochmans, Johan Neyts, Kai Dallmeier, Piet Maes, Dirk Daelemans

## Abstract

The ongoing COVID-19 pandemic is responsible for worldwide economic damage and nearly one million deaths. Potent drugs for the treatment of severe SARS-CoV-2 infections are not yet available. To identify host factors that support coronavirus infection, we performed genome-wide functional genetic screens with SARS-CoV-2 and the common cold virus HCoV-229E in non-transgenic human cells. These screens identified PI3K type 3 as a potential drug target against multiple coronaviruses. We discovered that the lysosomal protein TMEM106B is an important host factor for SARS-CoV-2 infection. Furthermore, we show that TMEM106B is required for replication in multiple human cell lines derived from liver and lung and is expressed in relevant cell types in the human airways. Our results identify new coronavirus host factors that may potentially serve as drug targets against SARS-CoV-2 or to quickly combat future zoonotic coronavirus outbreaks.

## INTRODUCTION

The recent pandemic of Severe Acute Respiratory Syndrome-Coronavirus-2 (SARS-CoV-2), the causative agent of COVID-19 (Coronavirus Disease 2019), has caused a worldwide health crisis with more than 33,000,000 confirmed infections and almost one million deaths, with numbers still rising (Dong et al., 2020). As for today, only two drugs have shown some clinical efficacy for the treatment of COVID-19 patients. The investigational antiviral drug remdesivir has been temporarily approved by authorities for emergency use on hospitalized patients with severe COVID-19, as it was shown in clinical trials to shorten the time to recovery in some cases (Beigel et al., 2020). Dexamethasone, an affordable and widely available steroid, can reduce mortality by one-third among patients critically ill with COVID-19, by supressing the hyperactive inflammatory immune response these patients suffer from in response to viral infection (Horby et al., 2020). Since both treatments only seem to benefit severe cases of COVID-19 and only to some limited extent, there is still an urgent need for efficient and safe therapeutic options to treat infected people while awaiting the development and worldwide implementation of safe and effective vaccines to halt the pandemic.

Coronaviruses are enveloped positive-sense RNA viruses that contain large genomes of up to 33.5 kb and have characteristic club-shaped spikes projecting from their virion surface (Fehr and Perlman, 2015). Coronaviruses cause respiratory and intestinal infections in a broad range of mammals and birds. Seven human coronaviruses (HCoVs) have been identified so far, which likely all emerged as zoonosis from bats, mice or domestic animals (Ye et al., 2020). The four so-called ‘common cold HCoVs’ are the alphacoronaviruses 229E and NL63, and betacoronaviruses OC43 and HKU1, which only cause mild upper respiratory tract illnesses (Liu et al., 2020). In contrast, the betacoronaviruses SARS-CoV, MERS-CoV and the recently emerged SARS-CoV-2, are highly pathogenic and can cause severe, potentially lethal respiratory infections. Since a large diversity of coronavirus types resides in animals and interspecies transmission frequently occurs (Chan et al., 2013; Cui et al., 2019; Ye et al., 2020), there is a high likelihood for the emergence of new pathogenic coronaviruses that can spread into the human population to pandemic proportions, as exemplified by the recent outbreak of SARS-CoV-2. Despite this risk and its great economic and social impact, our options to prevent or treat coronavirus infections remain very limited. Hence, the development of broad-spectrum antiviral drugs against members of this virus family could help not only to address the current high medical need, but also to quickly combat and contain zoonotic events in the future.

To develop such drugs, it is crucial to understand which host factors coronaviruses require to infect a cell, as in principle each step of the coronavirus replication cycle (receptor binding, endocytosis, fusion, translation of viral replication proteins and structural proteins, genome replication, virion assembly and release), may serve as target for antiviral intervention. While the viral entry step of coronaviruses has been relatively well characterized, the host-virus interplay in later steps of the viral life cycle remains largely elusive. For SARS-CoV-2, previous studies have shown that the protein angiotensin-converting enzyme 2 (ACE2) can serve as a receptor in Vero E6 cells (Wei et al., 2020) or in human cells overexpressing ACE2 (Hoffmann et al., 2020; Zhou et al., 2020; Zhu et al., 2020). In addition, it was shown that the SARS-CoV-2 spike (S) can be primed for fusion by cellular proteases such as furin, transmembrane serine protease 2 (TMPRSS2) and cathepsin B or L, depending on the target cell type (Hoffmann et al., 2020; Shang et al., 2020).

In this study, we performed a series of genome-wide CRISPR-based genetic screens in human cells to identify host factors required for SARS-CoV-2 and HCoV-229E infection. We identified PI3K type 3 as a common host factor for SARS-CoV-2, HCoV-229E, and HCoV-OC43, and show that small molecules targeting this protein might serve as broadly applicable anti-coronavirus inhibitors. Furthermore, we discovered that the lysosomal protein TMEM106B serves as an important specific host factor for SARS-CoV-2 infection in multiple liver- and lung-derived human cell lines.

## RESULTS

### Genome-wide knockout screens in human cells identify host factors for HCoV-229E and SARS-CoV-2

Genome-wide knockout screens have been widely used to identify host factors for various viruses (Flint et al., 2019; Li et al., 2020) but the only coronavirus for which genome-wide knockout screens have been performed to date is SARS-CoV-2. These screens were performed either in the African green monkey cell line Vero E6, or in human cells that were engineered to overexpress ACE2. Here, we chose to perform a CRISPR-based genome-wide knockout screen in human cells without introducing an exogenous receptor. To that end, we used the human liver cell line Huh7 in which our clinical isolate of SARS-CoV-2 was naturally able to induce a clear cytopathic effect (CPE). We performed screens with both SARS-CoV-2 as well as the less pathogenic HCoV-229E (**Figure 1A-D**). This allowed us to identify both (i) broad-spectrum coronavirus host factors as well as (ii) specific host factors for SARS-CoV-2 and HCoV-229E. Huh7 cells were transduced with the Brunello genome-wide library (Doench et al., 2016), treated with puromycin to eliminate untransduced cells, and then selected for survival during infection with either coronavirus, followed by sgRNA identification in the remaining cell population by deep sequencing for target deconvolution. We performed high stringency screens for HCoV-229E (**Figure 1B**) and SARS-CoV-2 (**Figure 1C**) by exposing cells to the virus until nearly all cells had died. For SARS-CoV-2, we also performed a lower stringency screen (**Figure 1D**) to identify genes with a more subtle effect on viral infection. However, low stringency screens may have an increased background and rather select for genes related to cell proliferation or general stress responses. The high stringency SARS-CoV-2 screen (**Figure 1C**) identified one significantly enriched gene, *TMEM106B*, while a larger number of genes was enriched in the low stringency screen (**Figure 1B)**. In contrast to previously published genome-wide screens with SARS-CoV-2, our screen did not identify *ACE2*. This difference is probably due to the fact that previous screens were performed in human cells that overexpress ACE2 (Heaton et al., 2020; Zhu et al., 2020) or in Vero E6 cells (Wei et al., 2020). Vero E6 cells express high levels of ACE2, whereas ACE2 is expressed at very low levels in Huh7 cells and several human airway cell lines (Clausen et al., 2020) (**Figure S1**). Yet, our screen with HCoV-229E identified *ANPEP* (**Figure 1C**), which encodes the well-known HCoV-229E receptor aminopeptidase N (AP-N). Comparison of the screens pointed towards *PIK3C3* and *TMEM41B* as common genes that were identified for both viruses, and we therefore selected these genes, as well as *TMEM106B*, for further validation.

**Figure 1.**
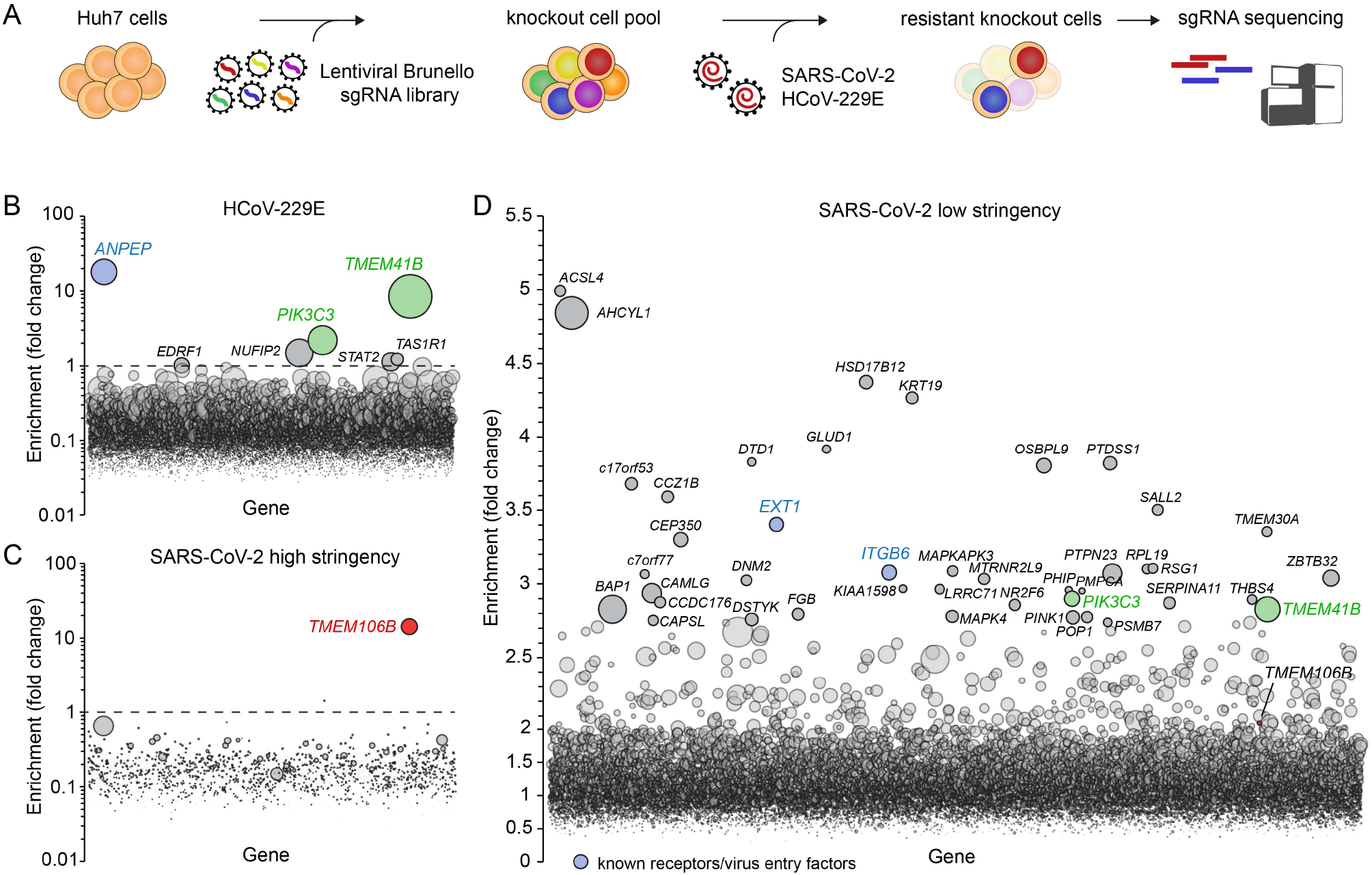
Genome-wide knockout screens in human cells identify host factors for SARS-CoV-2 and HCoV-229E infection. **A**) Overview of experimental steps performed during a genome-wide screen for coronavirus host factors. **B-D**) Genome-wide knockout screens were performed in Huh7 cells, with strong selection (high stringency) using HCoV-229E (B) and SARS-CoV-2 (C) or with mild SARS-CoV-2 selection (low stringency) (D). Each circle represents a gene, with size corresponding to significance of enrichment. The y-axis shows the enrichment of sgRNAs after virus selection compared to an uninfected control population (D) or the population on the first day of the screen prior to infection (B and C). Genes distributed on the x-axis in alphabetical order.

### PI3K type 3 is a druggable target against SARS-CoV-2 and other coronavirus infections

*PIK3C3* encodes PI3K type 3, the catalytic subunit of the PI3K complex that mediates the formation of phosphatidylinositol 3-phosphate and plays a role in many processes, including endocytic trafficking and the initiation and maturation of autophagosomes (Backer, 2016). Because *PIK3C3* is essential for cell survival (https://depmap.org/portal/depmap/), generation of PI3K type 3 null cells for genetical hit validation is not possible. We therefore confirmed the role of this factor in coronavirus infection instead by using the pharmacological inhibitors VPS34-IN1, VPS34-IN2, SAR405, and autophinib, which are structurally distinct inhibitors directly targeting PI3K type 3 (Bago et al., 2014; Pasquier et al., 2015; Robke et al., 2017; Ronan et al., 2014). As expected, all PI3K type 3 inhibitors inhibited the formation of LC3-positive autophagosome puncta and induced large vacuoles in treated cells (**Figure S2A**) (Ronan et al., 2014). PI3K type 3 inhibitors showed antiviral activity against SARS-CoV-2 in Vero E6 cells (**Figure 2A**) and Huh7 cells (**Figure 2B**) and were also active against HCoV-OC43 (**Figure 2C**) and the more distantly related alphacoronavirus HCoV-229E (**Figure 2D**). A time series experiment showed that inhibition of HCoV-229E by SAR405 occurs later in the viral life cycle than the attachment stage, as benchmarked by use of the attachment inhibitor UDA, and earlier than onset of intracellular replication as identified by use of remdesivir, an inhibitor of viral RNA synthesis (**Figure S2B**). This suggests a role of PI3K type 3 in an early step of the viral life cycle, but downstream of receptor binding.

**Figure 2.**
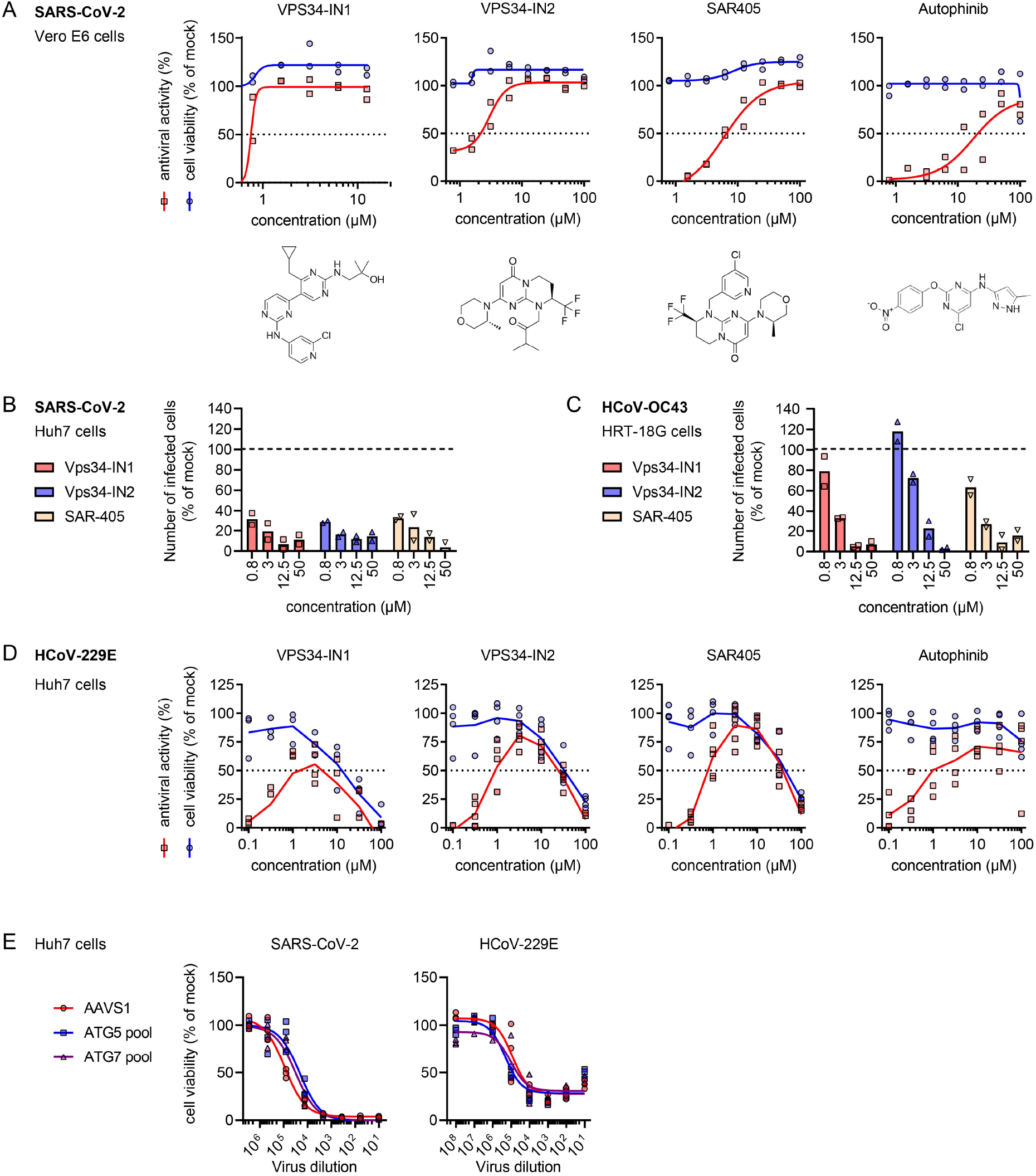
PI3K type 3 is a druggable target against SARS-CoV-2 and other coronaviruses. **A**) Vero E6 cells constitutively expressing EGFP were pretreated for 1 day with the indicated compounds and infected with SARS-CoV-2. Four days post infection, viability of cells was determined by measuring EGFP signal (scoring for surviving cells following infection) or by MTS assay (scoring for cell viability/general compound toxicity in uninfected control cells). Antiviral activities were calculated by subtracting the backgound (signal from infected untreated controls) and normalizing to uninfected untreated controls **B-C**) Number of infected cells after treatment with 12,5 μM of specific PI3K type 3 inhibitor for 6 hours, as compared to untreated control cells. The number of infected cells was quantified by high content image analysis after immunofluorescence staining for dsRNA. (B) Huh7 cells infected with SARS-CoV-2 (C) HRT-18G cells infected with HCoV-OC43. **D**) Huh7 cells were pretreated for 30 min with the indicated compounds and infected with HCoV-229E. At 3 days post infection, the cell viability was measured by MTS assay and plotted as a percentage of mock treated uninfected cells. **E**) Huh7 cells expressing pools of 4 sgRNAs targeting *ATG5, ATG7*, or the *AAVS1* safe targeting locus were infected with a dilution series of SARS-CoV-2 or HCoV-229E and incubated for three days at 35 °C, followed by measurement of the cell viability by MTS assay.

Since PI3K type 3 is involved in autophagosome formation, we investigated whether macroautophagy is required for SARS-CoV-2 and HCoV-229E infection. Huh7 cells expressing a pool of four sgRNAs targeting *ATG5* or *ATG7*, which are required for phagophore expansion (Dikic and Elazar, 2018), were unable to form LC3-positive autophagosomes (**Figure S2C**), indicating that the macroautophagy pathway was disrupted. However, *ATG5* and *ATG7* disruption did not affect the induction of CPE by SARS-CoV-2 or HCoV-229E (**Figure 2E**). Together, these results show that SARS-CoV-2 and other coronaviruses employ PI3K type 3 for infection, but do not depend on a functional macroautophagy pathway.

### Validation of TMEM106B, TMEM41B and EXT1 as SARS-CoV-2 or HCoV-229E host factors

To validate the findings from our genetic screens, we individually expressed sgRNAs targeting the identified genes, as well as the known receptor genes *ANPEP* and *ACE2*, in Huh7 cells and tested whether their abblation affected the sensitivity of cells to CPE induced by HCoV-229E or SARS-CoV-2. sgRNAs targeting *ANPEP* protected cells from HCoV-229E-induced cell death, as determined by crystal violet staining of surviving cells (**Figure 3A**). Cells were only partially protected from SARS-CoV-2-induced CPE by sgRNAs targeting *ACE2*, which is in line with the absence of *ACE2* in our screens. This limited effect of *ACE2* sgRNAs is probably due to the very low ACE2 expression in the human cell lines used, in contrast to human cell lines engineered to overexpress ACE2 or Vero E6 cells (**Figure S1**), in which others have found that SARS-CoV-2 infection depends on ACE2 (Wei et al., 2020).

**Figure 3.**
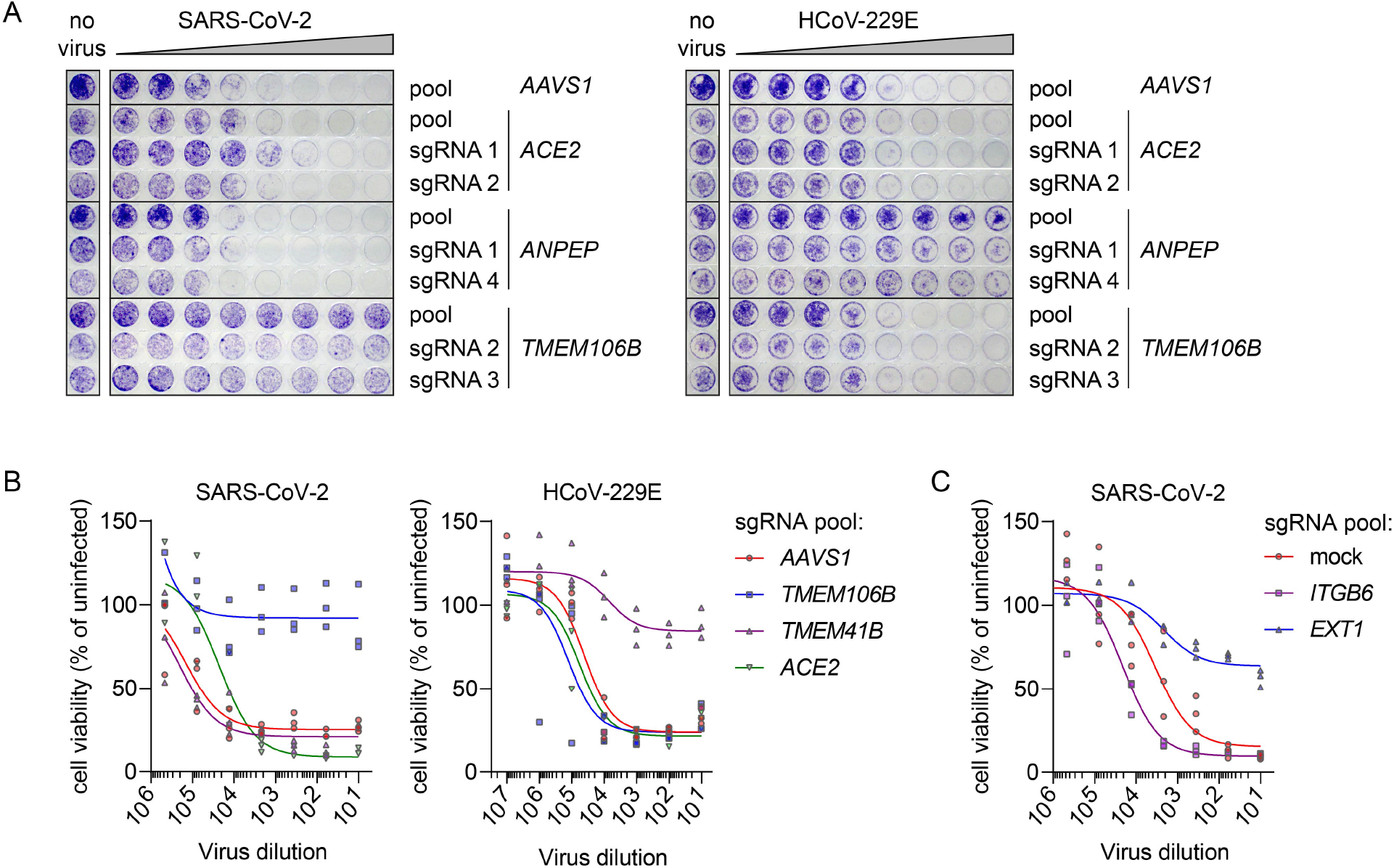
Validation of TMEM106B and EXT1 as SARS-CoV-2 host factors and TMEM41B as a HCoV-229E host factor. **A**) Huh7 cells expressing control sgRNAs (targeting the safe harbour gene *AAVS1)*, a gene-specific pool of 4 sgRNAs, or individual sgRNAs, were infected with a dilution series of SARS-CoV-2 (6-fold dilutions) or HCoV-229E (10-fold dilutions) and incubated for three days at 35 °C, followed by fixation and staining of surviving cells with crystal violet. **B**) Huh7 cells expressing the indicated sgRNA pools were infected with a dilution series of SARS-CoV-2 or HCoV-229E and incubated for three days at 35 °C, followed by measurement of cell viability by MTS assay. **C**) Huh7 cells expressing the indicated sgRNA pools were infected with a dilution series of SARS-CoV-2 and incubated for three days at 35 °C, followed by measurement of cell viability by MTS assay.

*TMEM106B* sgRNAs were highly enriched in our high stringency SARS-CoV-2 screen. This gene encodes the yet poorly understood protein TMEM106B, which is involved in lysosomal function and implicated in neurodegenerative disorders (Nicholson and Rademakers, 2016). sgRNAs targeting *TMEM106B* protected Huh7 cells from CPE caused by SARS-CoV-2, even when exposed to high virus concentrations, but had not effect on cells infected with HCoV-229E (**Figures 3A and B**). In contrast, sgRNAs targeting *TMEM41B*, which is involved in the early stage of autophagosome formation (Morita et al., 2018), only prevented CPE induced by HCoV-229E, but had no effect on SARS-CoV-2-infected cells in this assay (**Figure 3B**). This observation, and the fact that *TMEM41B* was identified only in the low stringency, but not the high stringency SARS-CoV-2 screen (**Figure 1C and D)**, suggests that SARS-CoV-2 only very weakly depends on *TMEM41B*.

Since *ACE2* was not identified in our screen, we questioned whether SARS-CoV-2 infection might be supported by other genes known to be involved in virus entry that were enriched in the SARS-CoV-2 low stringency screen; *ITGB6* and *EXT1* (**Figure 1B**). Both in wildtype Huh7 cells (**Figure S3**) and cells expressing *ACE2* sgRNAs, cells were protected against SARS-CoV-2 by sgRNAs targeting *EXT1*, which encodes exostosin-1 and is required for the synthesis of heparan sulfate. This observation is in line with a recent report showing that SARS-CoV-2 infection can be mediated by heparan sulfate (Clausen et al., 2020). Taken together, these results point towards a role of heparan sulfate in SARS-CoV-2 infection and identify TMEM41B and TMEM106B as specific host factors for HCoV-229E or SARS-CoV-2 infection, respectively.

### TMEM106B is required for SARS-CoV-2 infection in multiple human cell types

Our high stringency SARS-CoV-2 screen identified an important role of TMEM106B, a lysosomal protein that has never been implicated as a host factor for any pathogen before. To further corroborate the importance of TMEM106B for SARS-CoV-2 infection, we investigated its role in cell lines other than liver-derived Huh7 cells, including lung derived cell lines. Therefore, we examined several cell lines for susceptibility to SARS-CoV-2-induced CPE and selected the liver-derived cell lines Hep3B and lung-derived cell lines A549, NCI-H1975, and NCI-H2110. We next expressed *TMEM106B* sgRNAs in these cell lines. *TMEM106B* sgRNAs protected consistently against CPE caused by SARS-CoV-2 infection in Hep3B, A549, or NCI-H1975 cells, as determined by crystal violet staining (**Figure 4A**). Also NCI-H2110 cells, in which SARS-CoV-2 causes only a very limited CPE, were protected against SARS-CoV-2-induced cell death by sgRNAs targeting *TMEM106B*, as determined by MTS assay (**Figure 4B**). To further confirm the role of TMEM106B in SARS-CoV-2 infection, double-stranded RNA intermediates of viral RNA replication were visualized by immunofluorescence staining in infected Hep3B cells containing *TMEM106B* sgRNAs at 6 hours post infection. *TMEM106B* sgRNAs reduced the number of SARS-CoV-2-infected Hep3B cells as compared to cells containing control sgRNAs (**Figure 4C**). In addition, we also observed a reduction in virus progeny released in supernatant of infected NCI-H1975 cells containing *TMEM106B* sgRNAs as compared to control sgRNAs at 2 days post infection (**Figure 4D**). To investigate whether *TMEM106B* is present in relevant cell types in the human airways, we analyzed single cell sequencing data from COVID-19 patients (Chua et al., 2020). This showed that *TMEM106B* is expressed in ciliated and secretory cells, which are the main SARS-CoV-2 susceptible airway cell types (Chua et al., 2020; Zhu et al.). Together, these results indicate that TMEM106B is required for productive infection of human cell lines of different origins with SARS-CoV-2, and is expressed in relevant cell types in the human airways.

**Figure 4.**
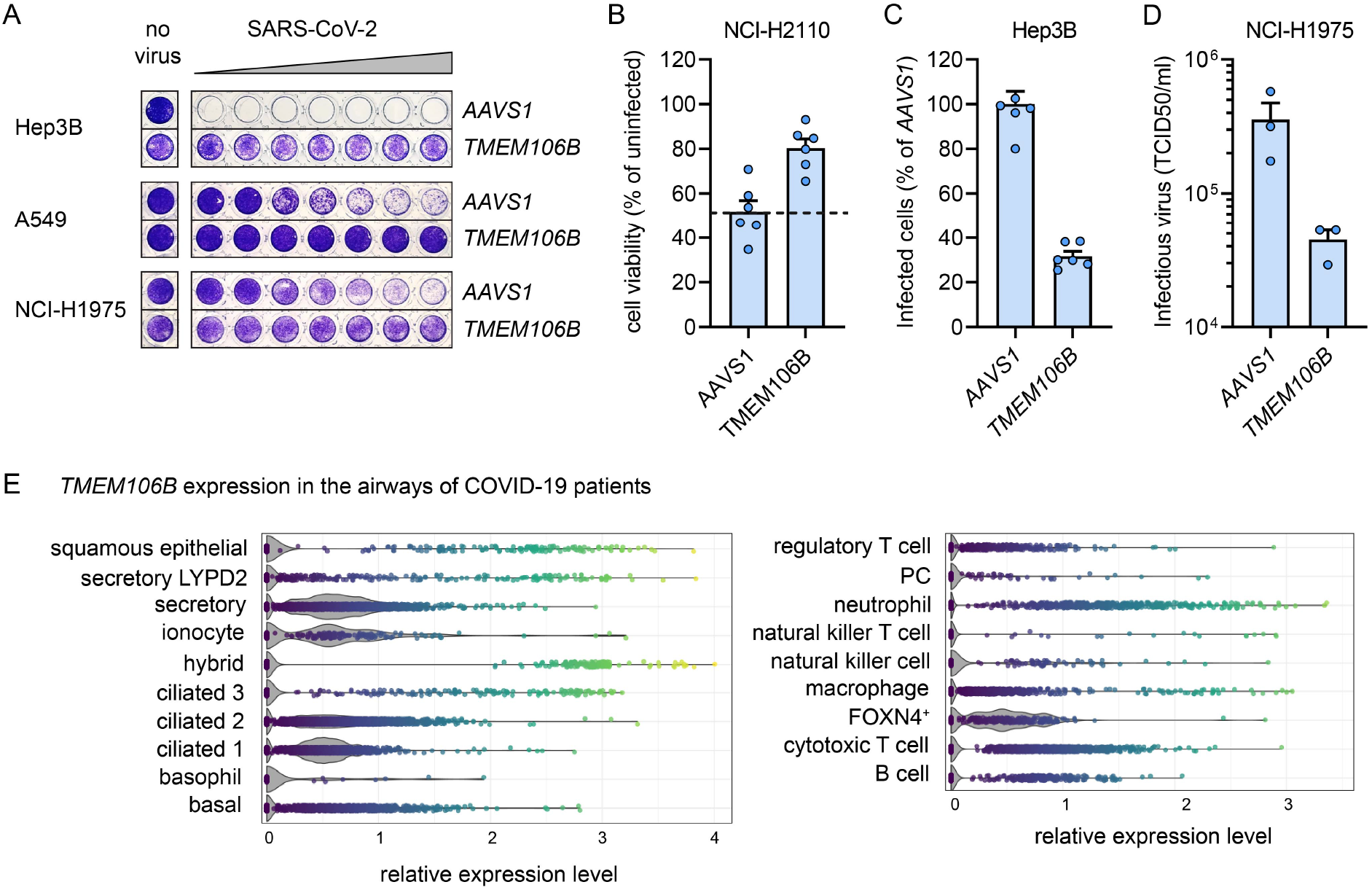
TMEM106B is required for SARS-CoV-2 infection in multiple human cell types and is expressed in human airways. **A**) Huh7 cells expressing control sgRNAs (targeting the safe harbour gene *AAVS1)* or a pool of 4 sgRNAs targeting *TMEM106B* were infected with 6-fold dilutions of SARS-CoV-2 and incubated 3 days (Hep3B) or 8 days (A549) at 35 °C or 6 days at 37 °C (NCI-H1975), followed by fixation and staining of intact cells with crystal violet. **B**) NCI-H2110 cells expressing pools of 4 sgRNAs were infected with SARS-CoV-2 at a MOI of ~0.2 and incubated for 7 days at 37 °C, followed by measurement of the cell viability by MTS assay. Triplicate data from two independent experiments are shown. **C**) Hep3B cells expressing pools of 4 sgRNAs were infected with a MOI of ~40 and stained for dsRNA at 6 hours post infection. The percentage of infected cells was determined by high content image analysis. Triplicate data from two independent experiments are shown. **D**) NCI-H1975 cells expressing pools of 4 sgRNAs were infected with SARS-CoV-2 at a MOI of ~1.5 and incubated for 2 days at 37 °C, after which the amount of infectious virus in the supernatant was determined by end-point-dilution on Vero E6 cells. Bars indicate the mean ±SEM. **E**) Analysis of single-cell sequencing data from two COVID-19 patients (Chua et al., 2020). Violin plots of *TMEM106B* expression levels are shown for single cells combined from nasopharyngeal swabs, bronchiolar protected specimen brushes, and bronchoalveolar lavages. Images were generated with the Magellan: COVID-19 Omics Explorer (https://digital.bihealth.org/).

## DISCUSSION

Although the receptor interactions and proteolytic activation of the SARS-CoV-2 S protein have been extensively studied, little is known about host factors required in following steps of viral infection. Here, we performed genome-wide genetic screens with SARS-CoV-2 and HCoV-229E and identified PI3K type 3 as a host factor shared by both viruses. Moreover, we show that the regulator of autophagy TMEM41B is required for HCoV-229E infection and identified TMEM106B as a new cellular host factor important for SARS-CoV-2 infection.

In our SARS-CoV-2 screens we did not identify *ACE2*, which is in contrast to previous screens performed in Vero E6 cells or engineered human cells overexpressing ACE2 (Heaton et al., 2020; Wei et al., 2020; Zhu et al., 2020). It has been reported that ACE2 overexpression in human cells enhances SARS-CoV-2 infection (Hoffmann et al., 2020; Zhou et al., 2020; Zhu et al., 2020) and that infection is inhibited by ACE2 depletion in Vero E6 cells (Wei et al., 2020). However, to our knowledge, it has not been investigated whether knockout of ACE2 in non-transgenic human cells prevents SARS-CoV-2 infection. We observed that all human cell lines tested in this study were susceptible to SARS-CoV-2 infection, despite very low levels of ACE2 expression. In addition, we found that heparan sulfate is important for efficient infection, both in wt Huh7 cells and cells expressing *ACE2* sgRNAs (**Figures 3 and S3**), which is in line with a recent study showing that heparan sulfate facilitates SARS-CoV-2 binding to cells (Clausen et al., 2020). Moreover, another recent study showed that two other human receptor proteins, KREMEN1 and ASGR1, can facilitate infection of SARS-CoV-2 S pseudotyped virus (Gu et al., 2020). Therefore, it is plausible that SARS-CoV-2, alike other viruses, has a broader repertoire of possible (redundant) cellular receptor than initially postulated.

PI3K type 3 inhibitors have antiviral activity against SARS-CoV-2, HCoV-229E, and HCoV-OC43, and may therefore have potential to serve as pan-coronavirus inhibitors. Our data (**Figure S2B**) suggest that PI3K type 3 plays a role in an early step of the viral life cycle, such as endocytosis, fusion, translation or early onset of replication. As recently demonstrated by others, treatment of SARS-CoV-2 infected cells with PI3K type 3 inhibitors causes dispersal of the viral N protein and dsRNA throughout the cytoplasm, suggesting a role of this factor in replication complex formation (Silvas et al., 2020). We showed that disruption of the autophagy genes *ATG5* and *ATG7*, which are required for phagophore expansion, does not negatively impact SARS-CoV-2 and HCoV-229E infection (**Figure 2E**). Thus, it is possible that PI3K type 3 supports infection by inducing phagophore nucleation, while the later stages of macroautophagy are not required.

Importantly, we identified a novel host factor required specifically for SARS-CoV-2 infection. TMEM106B is a 274 amino acid transmembrane protein that resides in endosomes and lysosomes, and controls lysosome size, number, mobility and trafficking. TMEM106B is not well characterized and only recently received attention because of its role in frontotemporal dementia, the second leading cause of pre-senile neurodegeneration after Alzheimer’s disease (Nicholson and Rademakers, 2016). TMEM106B plays a pivotal role in lysosomal acidification, via direct interaction with the proton pump vacuolar-ATPase accessory protein 1 (AP1) (Klein et al., 2017). Therefore, a possible role of TMEM106B might be to promote acidification of vesicles in the endolysosomal pathway, in order to facilitate efficient delivery of the SARS-CoV-2 genome into the cytoplasm. This is in agreement with our finding that TMEM106B plays a role early in the viral replication cycle, i.e. within the first 6 hours after infection (**Figure 4C**) and with recent findings that entry of SARS-CoV-2 S pseudotyped virus depends on endosomal acidification (Hoffmann et al., 2020; Ou et al., 2020). Interestingly, HCoV-229E also requires endosomal acidification (Blau and Holmes, 2001), but does not require TMEM106B for infection. This suggests that different, though related, viruses may depend on distinct factors to exploit similar cellular pathways. Alternatively, TMEM106B may function as a lysosomal receptor for SARS-CoV-2, similar to the lysosomal receptor NPC1 used by Ebola virus (Carette et al., 2011). Further studies are needed to precisely establish which stage of infection is supported by TMEM106B.

As new pathogenic coronaviruses periodically emerge, these viruses will continue to pose a public health threat beyond the ongoing COVID-19 pandemic, warranting the development of potent coronavirus inhibitors. Here, we used a genome-wide knockout approach to identify coronavirus host factors in human cells and discovered TMEM106B as a potential new target that might be exploited in the development of drugs to counter the current pandemic or future outbreaks of pathogenic coronaviruses.

## ACKNOWLEDGEMENTS

We thank Niels Willems, Nathalie van Winkel, Liesbeth Mercelis, Kristien Minner, Lotte Bral, and Bob Massant for exceptional technical help and support. S.J., J.N. and K.D. were supported by the Research Foundation Flanders (FWO) under the Excellence of Science (EOS) program (VirEOS project 30981113).

## AUTHOR CONTRIBUTIONS

J.B., S.J. and P.M. performed genetic screens. J.B., L.P., S.J. performed infectivity assays. L.P., E.V., and M.J. performed other experiments. J.B., L.P., E.V., and M.J. were involved in data analysis. J.B., L.P., E.V., M.J., and D.D designed the project. D.J., J.N., K.D., P.M., and D.D. supervised and supported the project. J.B., L.P., E.V., M.J., and D.D. co-wrote the manuscript.

## METHODS

### Chemicals and reagents

Reference inhibitor compounds Autophinib and VPS34-IN1 were purchased from Selleckchem and SAR405 and VPS34-IN2 were obtained from MedChemExpress. Plant lectin Urtica dioica agglutinin (UDA), isolated from the Urtica dioica rhizomes, was kindly donated by E. Van Damme (Ghent, Belgium). Chloroquine was purchased from Acros Organics and Remdesivir was ordered from MedKoo. Stock solutions were prepared in DMSO.

### Cell culture

HEK293T (received from prof. Jason Moffat, Donnelly Centre, University of Toronto, Canada), Vero E6, Huh-7 (CLS - 300156; human hepatoblastoma), Hep3B (ATCC HB-8064; human hepatocellular carcinoma) and HRT-18G (ATCC CRL-11663; human colorectal adenocarcinoma) were maintained in Dulbecco’s Modified Eagle Medium (DMEM, Gibco Life Technologies) supplemented with 8% heat-inactivated fetal bovine serum (HyClone, GE Healthcare Life Sciences), 0.075% sodium bicarbonate (Gibco Life Technologies) and 1mM sodium pyruvate (Gibco Life Technologies). A549 cells were maintained in F-12K medium supplemented with 10% heat-inactivated fetal bovine serum. NCI-H1975 (ATCC-CRL-5908) and NCI-H2110 (ATCC-CRL-5924) cells were maintained in RPMI medium supplemented with 10% heat-inactivated fetal bovine serum. All cell lines were maintained at 37°C under 5% CO2.

### Generation of virus stocks

SARS-CoV-2 strain BetaCov/Belgium/GHB-03021/2020 (EPI ISL 407976|2020-02-03) was recovered from a nasopharyngeal swab taken from a RT-qPCR-confirmed asymptomatic patient returning from Wuhan in February 2020. Infectious virus was isolated and multiplied by five serial passages on Huh7 cells. Cells were seeded in DMEM supplemented with 10% heat-inactivated fetal bovine serum to reach a confluency of ~80% the next day. After replacing the medium by DMEM + 2% or 4% fetal bovine serum, cells were infected with SARS-CoV-2 at a MOI of ~0.01. When most cells were dying, supernatant was removed from the cells, centrifuged to remove cell debris and stored at −80 °C. The HCoV-229E (ATCC VR-740) and HCoV-OC43 (ATCC VR-1558) virus stocks were obtained by inoculating a confluent monolayer of Huh7 or HRT-18G cells, respectively. The supernatant was harvested after 3 days of incubation for HCoV-229E, or 7 days of incubation for HCoV-OC43, at 35 °C under 5% CO2 and stored in aliquots at −80°C, after one freeze-thaw cycle and removal of cellular debris by centrifugation.

### Genome-wide knockout screens

For the HCoV-229E and SARS-CoV-2 (high stringency) screens, 1.5 x 10^8^ Huh7 cells for each of two replicates were transduced at a MOI of ~0.3 with lentivirus containing the Brunello genome-wide library in lentiCRISPRv2 (Addgene 73179), which contains 77.441 sgRNAs targeting 19.114 genes. Cells were selected with 2 μg/ml puromycin for three days to eliminate untransduced cells, seeded at a coverage of ~200 cells/sgRNA for each replicate and infected with HCoV-229E or SARS-CoV-2. Surviving cells were harvested at 18 days post infection (HCoV-229E) or 41 days post infection (SARS-CoV-2). For the SARS-CoV-2 low stringency screen, 1.5 x 10^8^ Huh7 cells for each of two replicates were transduced at a MOI of ~0.2 with lentivirus containing the Brunello library and selected with puromycin for three days. Then, cells were seeded at a coverage of 500 cells/sgRNA for each replicate in DMEM with 4% fetal bovine serum and infected with SARS-CoV-2 at a MOI of 0.1. For each replicate, uninfected cells were maintained under similar conditions as the infected cells and harvested simultaneously. Five days post infection, cells were cultivated in DMEM with 20% fetal bovine serum for three days to allow cell recovery, and were infected again with a MOI of ~0.1. Cells were harvested at 14 days post infection. Genomic DNA was extracted from cells using the QIAmp DNA Mini Kit (Qiagen ref. 51306) or, for the SARS-CoV-2 low stringency screen with the QIAmp DNA Blood maxi kit (Qiagen ref. 51194). In a first PCR step, regions of ~600 bp containing the sgRNA sequence were amplified using NEBNext Ultra II Q5 Master Mix (NEB #M0544S) in 25 amplification cycles. A second PCR of 10 cycles with NEBNext Ultra II Q5 Master Mix was performed with primers containing Illumina adapters and TruSeq indexes. Products were separated by agarose gel electrophoresis and purified with the PureLink Quick Gel Extraction Kit (Thermo Fisher K210012). Samples were then diluted to 2–4 nM, pooled, and denatured and diluted according to the instructions for single-end sequencing on a MiSeq (Illumina) with a MiSeq-v2 50 cycles or a MiSeq-v3-150 cycles kit (Illumina) and 10% PhiX (Illumina) spike-in. FastQ files were further analyzed with CRISPRCloud2 (Jeong et al., 2018).

### Genome-wide knockout screen hit validation via cell viability assays

For individual validation of genes, guides enriched during the genome-wide knockout screens were cloned into the pLentiCRISPRv2 plasmid (Addgene 52961) following the standard cloning protocol. For lentiviral particle production, HEK293T cells were plated in 40 mL supplemented DMEM in T150 (TPP) flasks at 45% confluency and incubated overnight. 24 hours later, the cells were transfected using X-TremeGENE 9 (Roche) with the pLentiCRISPR plasmids and the lentiviral packaging plasmids pMD2.G and psPAX2 to generate lentiviral particles coated with the VSV-G protein and incubated overnight. 24 hours post transfection the medium was changed to DMEM supplemented with serum-free BSA growth media (DMEM + 1.1g/100mL BSA and 20 μg/mL gentamicin). The supernatant containing lentiviral particles was harvested 72 hours after transfection and stored at −80 °C. Cells were transduced with lentiviruses expressing only one sgRNA or a pool of the 4 sgRNAs from the Brunello genome-wide knockout library and then selected with puromycin for 3 days (sgRNAs target sequences are in **Table S1**). Cells stably expressing specific sgRNAs were seeded in 96-well plates at 4000 cells/well in medium with 8% or 10% fetal bovine serum. The following day, serial dilutions of virus in medium without fetal bovine serum were added to the cells, resulting in a serum concentration of 4% or 5%. Cells were incubated until sufficient CPE was visible. For MTS assays, medium was removed from the cells and replaced by MTS reagent (CellTiter 96 AQueous One Solution Cell Proliferation Assay from Promega, Madison, WI) diluted in PBS. The absorbance was measured with a Tecan Spark microplate reader. For crystal violet staining, cells were fixed in 4% formaldehyde for 30 min, stained with a 1% crystal violet solution in water, and rinsed with water.

### Virus inhibition assays

The antiviral activity of PI3K type 3 inhibitors on SARS-CoV-2 in Vero E6 cells was evaluated as follows: on day −1, the test compounds were serially diluted in DMEM (Gibco cat no 41965-039) supplemented with 2% v/v heat-inactivated FCS and sodium bicarbonate (Gibco 25080-060). Diluted compounds were then mixed with EGFP-expressing Vero E6 cells at 25,000 cells/well in 96-well plates (Greiner Bio-One, Vilvoorde, Belgium; Catalog 655090). The plates were incubated overnight in a humidified incubator at 37°C and 5% CO2. On day 0, SARS-CoV-2 was added at 20 TCID50/well and on day 4 post infection, the wells were examined for EGFP expression using a high-content imaging platform and the images of the wells were converted into signal values. To obtain values for antiviral activity, the background signal was subtracted based on infected-untreated controls and signal values were normalized to uninfected-untreated controls. Toxicity of compounds in the absence of virus was evaluated by MTS assay. All compounds were tested in duplicate, in two independent experiments. To evaluate the antiviral activity of PI3K type 3 inhibitors against HCoV-229E, Huh7 cells were seeded into 384-well plates. The next day, serial dilutions of the compounds were added to the cells prior to infection with HCoV-229E at 30 TCID50 (50% tissue culture infective doses) per well. At 3 days post infection, the virus-induced CPE was measured by MTS assay. The compounds were tested in at least four independent experiments.

### Time of drug addition assay

Huh-7 cells were seeded into 48-well dishes at 40,000 cells per well. After 24 hours of incubation at 37°C, the cells were cooled on ice for 1 hour, followed by addition of 30 CCID50 of the HCoV-229E virus and further incubation at 35°C. The test compounds were added at a concentration approximately 10-fold above their EC50, at different time points post infection (−30 min, 0h, 30 min, 1h, 2h, 3h, 5h and 8h p.i.). At 11 h p.i., total cellular RNA extracts were prepared and viral RNA was quantified using the CellsDirect One-Step qRT-PCR kit (Thermo Fisher Scientific). One-step real-time RT-PCR was performed using the 229E-FP forward primer (5’-TTCCGACGTGCTCGAACTTT-3’), 229E-RP reverse primer (5’-CCAACACGGTTGTGACAGTGA-3’) and the TaqMan minor groove binder (MGB) probe 229E-TP (FAM-5’-TCCTGAGGTCAATGCA-3’-NFQ-MGB; Thermo Fisher Scientific), derived from the HCoV 229E membrane protein gene sequence as described previously (Vijgen et al., 2005). Amplification and detection were performed in an ABI 7500 Fast Sequence Detection System (Applied Biosystems, Foster City, CA, USA) under the following conditions: an initial reverse transcription at 50 °C for 15 min, followed by PCR activation at 95 °C for 2 min and 45 cycles of amplification (15 s at 95 °C and 30 s at 60 °C). Six independent experiments were carried out.

### Immunofluorescence assays

Immunofluorescence staining was performed according to standard procedures. Briefly, all cells were seeded at a density of 20.000 cells per well in 8-well μ-slides (Ibidi). Cells were allowed to adhere overnight before receiving compound treatment and/or viral infection with SARS-CoV-2 or HCoV-OC43. After incubation, cells were fixed (4% PFA in PBS), washed and permeabilized (0.2% Triton X-100 in PBS). Employed primary antibodies were rabbit anti-LC3B (L7543, Sigma) at a 1:200 dilution and mouse anti-dsRNA (J2, Scicons) at a 1:1000 dilution. Secondary antibodies Alexa Fluor® 568 goat anti-rabbit (A11011, Invitrogen, ThermoFisher Scientific) and Alexa Fluor® 488 goat anti-mouse (A11029, Invitrogen, ThermoFisher Scientific) were diluted 1:500. Cell nuclei were counterstained with DAPI and the samples were imaged by confocal microscopy on a Leica TCS SP5 confocal microscope (Leica Microsystems), employing a HCX PL APO 63x (NA 1.2) water immersion objective. The percentage of infected cells was quantified by high content image analysis (ArrayScan XTI, ThermoFisher Scientific) for at least 3000 cells per condition.

### Generation of ACE2 overexpressing Huh7 cells

The pLCKO plasmid was a gift from Jason Moffat (Addgene plasmid #73311). The invariant gRNA scaffold was removed together with the puromycin resistance gene and replaced with *ACE2* (Addgene Plasmid #1786) followed by a P2A-coupled blasticidin resistance gene driven by a cytomegalovirus promotor. The resulting pLCKO-ACE2-P2A-Blasticidin vector was used to make lentiviral particles, as described above. Huh7 cells were transduced with a the lentiviral stock in the presence of polybrene (8 μg/ml). After 24 hours medium was replaced with medium containing Blasticidin (10 ug/ml) and cells were incubated for an additional 48 hours.

### Simple Western analysis

For Simple Western analysis, cells were lysed in RIPA lysis buffer (Sigma) for 1 hour at 4 °C. Whole cell lysates were cleared by centrifugation. Proteins were separated by size (12-230 kDa) and visualized on a Wes system (ProteinSimple, San Jose, CA, USA) with an anti-mouse or anti-goat IgG-HRP antibody (R&D systems, HAF109) detecting the primary antibody against GAPDH (Santa Cruz Biotechnology, sc-47724) or anti-hACE2 (R&D systems, AF933), respectively. Protein signals were visualized and quantified with the Compass software, v4.0.0 (Protein Simple).

## SUPPLEMENTAL INFORMATION

**Figure S1.**
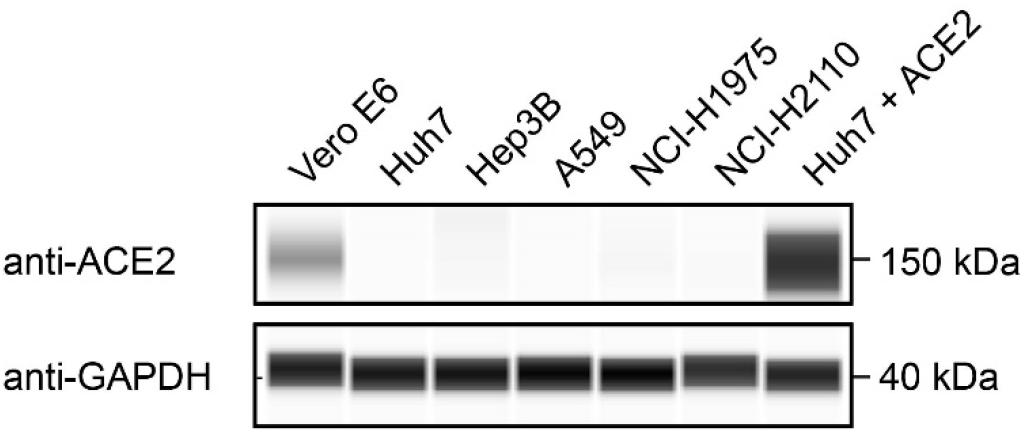
Analysis of ACE2 expression levels in different cell lines. Lysates of the indicated wildtype cell lines, or Huh7 cells transduced with an *ACE2* overexpression construct, were analyzed with a ProteinSimple Wes™ system, using antibodies specific for ACE2 and the endogenous control GAPDH.

**Figure S2.**
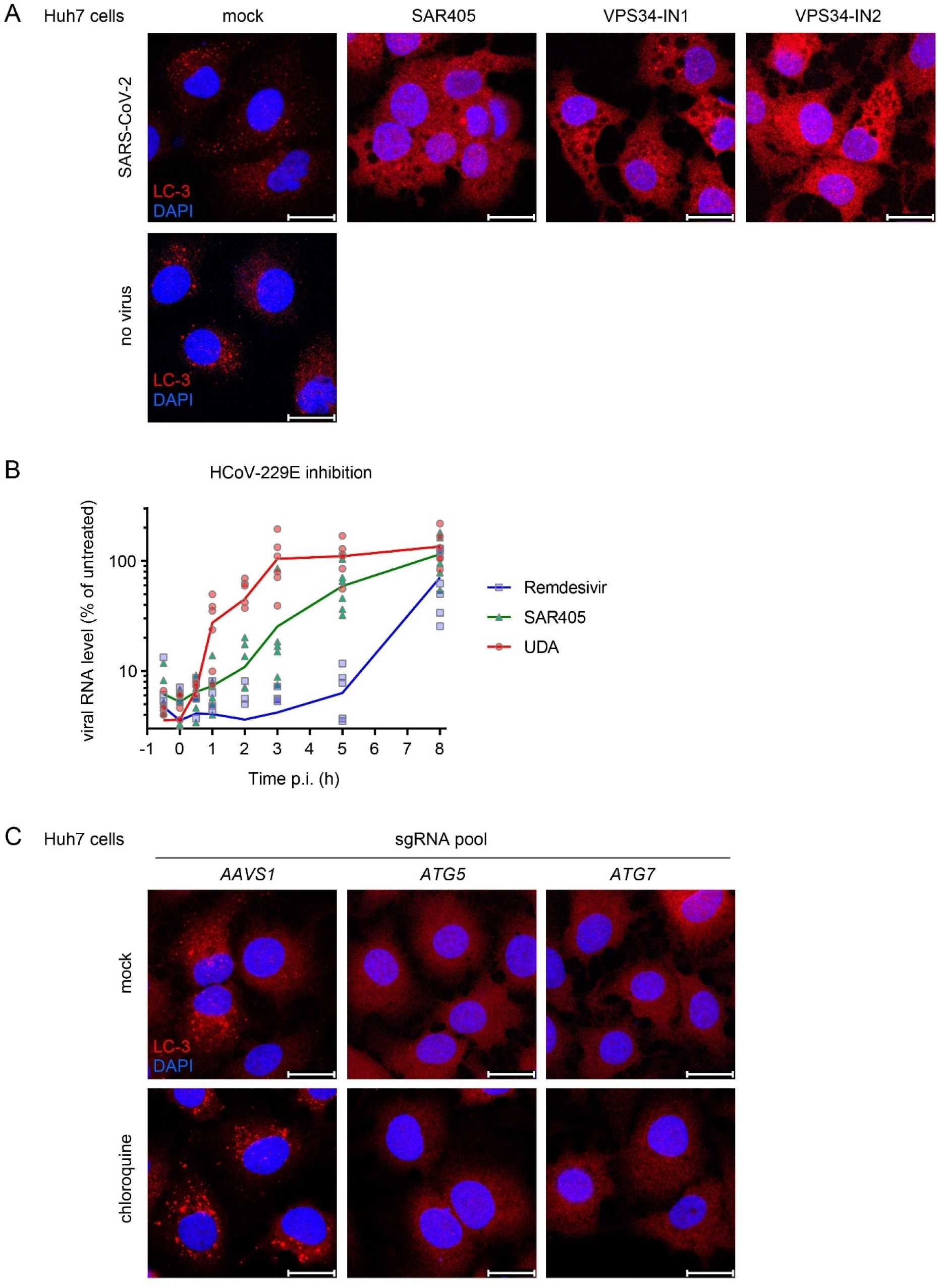
Coronavirus infection requires PI3K type 3 in an early step of the life cycle and does not require a functional macroauthophagy pathway. **A**) Immunofluorescence staining of LC3B in uninfected or SARS-CoV-2-infected Huh7 cells treated with 12.5 μM of specific PI3K type 3 inhibitors for 6 hours; PI3K type 3 inhibition completely abolishes the formation of LC3-positive puncta and induces vacuoles in treated cells. **B**) Time series experiment showing early stage post-receptor binding effect of PI3K type 3 inhibitor SAR405 on HCoV-229E infection. Huh7 cells were infected, treated with SAR405 at different timepoints, followed by determination of viral RNA levels at 10 hours post infection by qPCR. UDA: Urtica dioica agglutinin. Combined results of six independent experiments are shown **C**) Immunofluorescence staining of LC3B in Huh7 cells expressing a pool of four sgRNAs targeting *ATG5, ATG7*, or the *AAVS1* safe targeting locus. Chloroquine, an inhibitor of autophagic flux that decreases autophagosome-lysosome fusion, induces an increase in LC3-positive puncta in control cells, but fails to do so in *ATG5* and *ATG7* knockout cells, confirming the effective knockout of both genes (bar: 25 μm).

**Figure S3.**
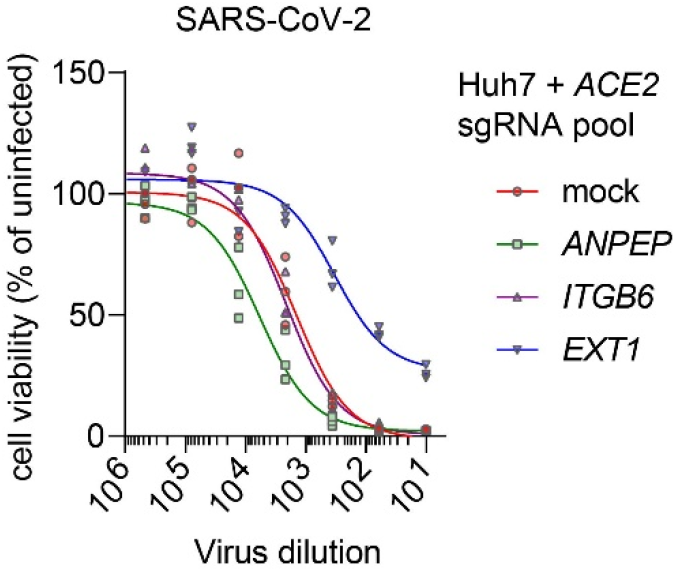
SARS-CoV-2 infection requires the heparan sulfate biosynthesis factor *EXT1*. Huh7 cells expressing a pool of sgRNAs targeting *ACE2*, together with the indicated sgRNA pools were infected with a dilution series of SARS-CoV-2 and incubated for three days at 35 °C, followed by measurement of cell viability by MTS assay.

**Table S1.**
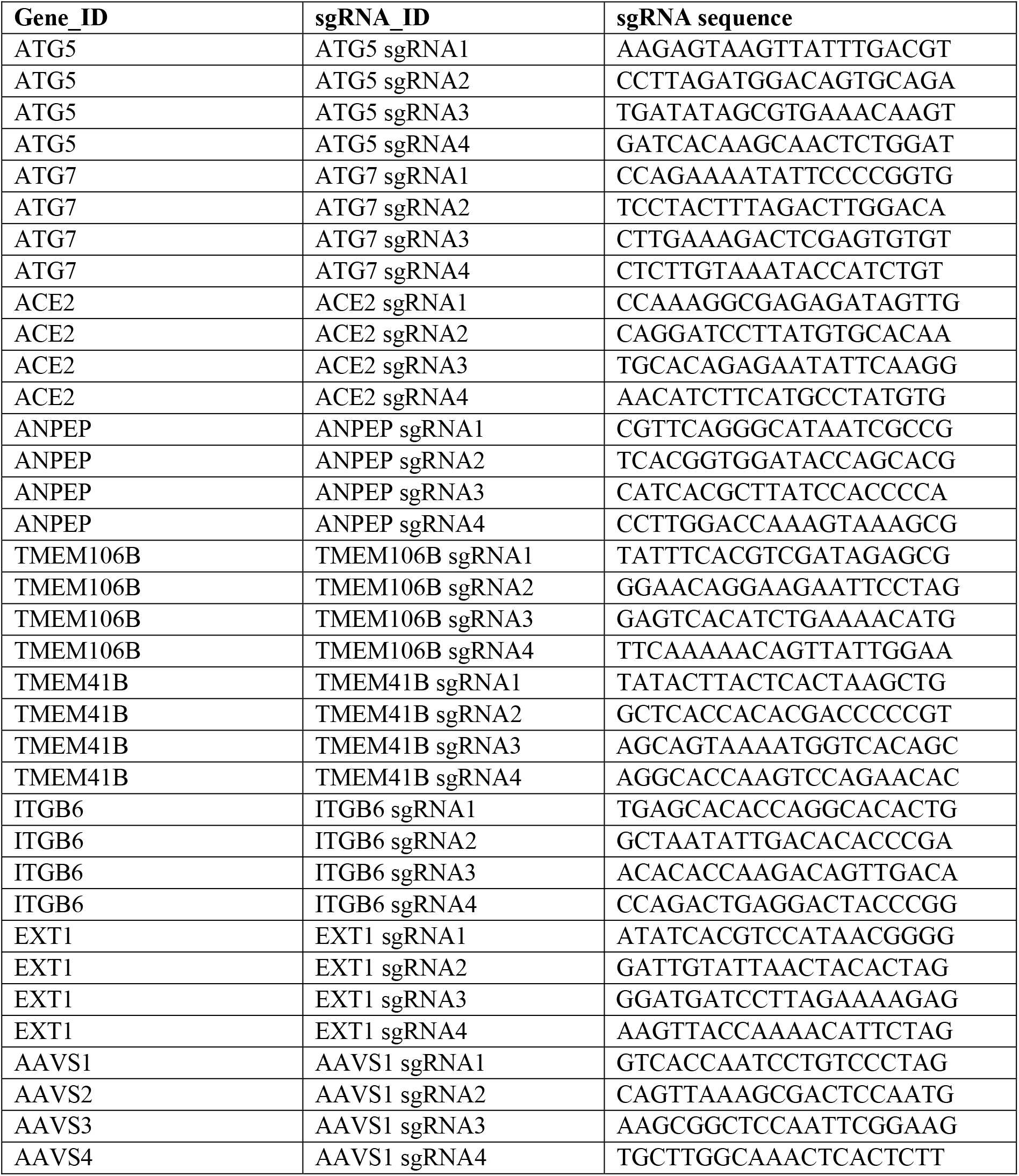
Sequences of sgRNAs used in this study

## REFERENCES

Backer, J.M. (2016). The intricate regulation and complex functions of the Class III phosphoinositide 3-kinase Vps34. Biochem. J. 473, 2251–2271.

Bago, R., Malik, N., Munson, M.J., Prescott, A.R., Davies, P., Sommer, E., Shpiro, N., Ward, R., Cross, D., Ganley, I.G., et al. (2014). Characterization of VPS34-IN1, a selective inhibitor of Vps34, reveals that the phosphatidylinositol 3-phosphate-binding SGK3 protein kinase is a downstream target of class III phosphoinositide 3-kinase. Biochem. J. 463, 413–427.

Beigel, J.H., Tomashek, K.M., Dodd, L.E., Mehta, A.K., Zingman, B.S., Kalil, A.C., Hohmann, E., Chu, H.Y., Luetkemeyer, A., Kline, S., et al. (2020). Remdesivir for the Treatment of Covid-19 — Preliminary Report. N. Engl. J. Med.

Blau, D.M., and Holmes, K. V (2001). Human coronavirus HCoV-229E enters susceptible cells via the endocytic pathway. Adv. Exp. Med. Biol. 494, 193–198.

Carette, J.E., Raaben, M., Wong, A.C., Herbert, A.S., Obernosterer, G., Mulherkar, N., Kuehne, A.I., Kranzusch, P.J., Griffin, A.M., Ruthel, G., et al. (2011). Ebola virus entry requires the cholesterol transporter Niemann-Pick C1. Nature 477, 340–343.

Chan, J.F., To, K.K., Tse, H., Jin, D.Y., and Yuen, K.Y. (2013). Interspecies Transmission and Emergence of Novel Viruses: Lessons From Bats and Birds. Trends Microbiol. 21.

Chua, R.L., Lukassen, S., Trump, S., Hennig, B.P., Wendisch, D., Pott, F., Debnath, O., Thürmann, L., Kurth, F., Völker, M.T., et al. (2020). COVID-19 severity correlates with airway epithelium–immune cell interactions identified by single-cell analysis. Nat. Biotechnol. 38.

Clausen, T.M., Sandoval, D.R., Spliid, C.B., Pihl, J., Painter, C.D., Thacker, B.E., Glass, C.A., Narayanan, A., Majowicz, S.A., Zhang, Y., et al. (2020). SARS-CoV-2 Infection Depends on Cellular Heparan Sulfate and ACE2. Cell 183.

Cui, J., Li, F., and Shi, Z.L. (2019). Origin and evolution of pathogenic coronaviruses. Nat. Rev. Microbiol. 17, 181–192.

Dikic, I., and Elazar, Z. (2018). Mechanism and medical implications of mammalian autophagy. Nat. Rev. Mol. Cell Biol. 19, 349–364.

Doench, J.G., Fusi, N., Sullender, M., Hegde, M., Vaimberg, E.W., Donovan, K.F., Smith, I., Tothova, Z., Wilen, C., Orchard, R., et al. (2016). Optimized sgRNA design to maximize activity and minimize off-target effects of CRISPR-Cas9. Nat. Biotechnol. 34, 184–191.

Dong, E., Du, H., and Gardner, L. (2020). An Interactive Web-Based Dashboard to Track COVID-19 in Real Time. Lancet. Infect. Dis. 20.

Fehr, A.R., and Perlman, S. (2015). Coronaviruses: An Overview of Their Replication and Pathogenesis. In Coronaviruses: Methods and Protocols, pp. 1–282.

Flint, M., Chatterjee, P., Lin, D.L., McMullan, L.K., Shrivastava-Ranjan, P., Bergeron, É., Lo, M.K., Welch, S.R., Nichol, S.T., Tai, A.W., et al. (2019). A genome-wide CRISPR screen identifies N-acetylglucosamine-1-phosphate transferase as a potential antiviral target for Ebola virus. Nat. Commun. 10, 1–13.

Gu, Y., Cao, J., Zhang, X., Gao, H., Wang, Y., Zhang, J., Shen, G., Jiang, X., Yang, J., Zheng, X., et al. (2020). Interaction network of SARS-CoV-2 with host receptome through spike protein. bioRxiv.

Heaton, B.E., Trimarco, J.D., Hamele, C.E., Harding, A.T., Tata, A., Zhu, X., Tata, P.R., Smith, C.M., and Heaton, N.S. (2020). SRSF protein kinases 1 and 2 are essential host factors for human coronaviruses including SARS-CoV-2. bioRxiv Prepr. Serv. Biol. 1–28.

Hoffmann, M., Kleine-Weber, H., Schroeder, S., Krüger, N., Herrler, T., Erichsen, S., Schiergens, T.S., Herrler, G., Wu, N.H., Nitsche, A., et al. (2020). SARS-CoV-2 Cell Entry Depends on ACE2 and TMPRSS2 and Is Blocked by a Clinically Proven Protease Inhibitor. Cell 1–10.

Horby, P., Lim, W.S., Emberson, J., Mafham, M., Bell, J., Linsell, L., Staplin, N., Brightling, C., Ustianowski, A., Elmahi, E., et al. (2020). Effect of Dexamethasone in Hospitalized Patients with COVID-19: Preliminary Report. N. Engl. J. Med.

Jeong, H., Kim, S.Y., Rousseaux, M.W.C., Zoghbi, H.Y., and Liu, Z. (2018). CRISPRCloud2: A cloud-based platform for deconvoluting CRISPR screen data. bioRxiv.

Klein, Z.A., Takahashi, H., Ma, M., Stagi, M., Zhou, M., Lam, T.T., Strittmatter, S.M., Klein, Z.A., Takahashi, H., Ma, M., et al. (2017). Loss of TMEM106B Ameliorates Lysosomal and Frontotemporal Dementia-Related Phenotypes in Article Loss of TMEM106B Ameliorates Lysosomal and Frontotemporal Dementia-Related Phenotypes in Progranulin-Deficient Mice. Neuron 95, 281–296.e6.

Li, B., Clohisey, S.M., Chia, B.S., Wang, B., Cui, A., Eisenhaure, T., Schweitzer, L.D., Hoover, P., Parkinson, N.J., Nachshon, A., et al. (2020). Genome-wide CRISPR screen identifies host dependency factors for influenza A virus infection. Nat. Commun. 11.

Liu, D.X., Liang, J.Q., and Fung, T.S. (2020). Human Coronavirus-229E, -OC43, -NL63, and -HKU1. Ref. Modul. Life Sci.

Morita, K., Hama, Y., Izume, T., Tamura, N., Ueno, T., Yamashita, Y., Sakamaki, Y., Mimura, K., Morishita, H., Shihoya, W., et al. (2018). Genome-wide CRISPR screen identifies TMEM41B as a gene required for autophagosome formation. J. Cell Biol. 217, 3817–3828.

Nicholson, A.M., and Rademakers, R. (2016). What we know about TMEM106B in neurodegeneration. Acta Neuropathol. 132, 639–651.

Ou, X., Liu, Y., Lei, X., Li, P., Mi, D., Ren, L., Guo, L., Guo, R., Chen, T., Hu, J., et al. (2020). Characterization of spike glycoprotein of SARS-CoV-2 on virus entry and its immune cross-reactivity with SARS-CoV. Nat. Commun. 11.

Pasquier, B., El-Ahmad, Y., Filoche-Rommé, B., Dureuil, C., Fassy, F., Abecassis, P.Y., Mathieu, M., Bertrand, T., Benard, T., Barrière, C., et al. (2015). Discovery of (2 S)-8-[(3 R)-3-Methylmorpholin-4-yl]-1-(3-methyl-2-oxobutyl)-2-(trifluoromethyl)-3,4-dihydro-2 H-pyrimido[1,2-a]pyrimidin-6-one: A novel potent and selective inhibitor of Vps34 for the treatment of solid tumors. J. Med. Chem. 58, 376–400.

Robke, L., Laraia, L., Carnero Corrales, M.A., Konstantinidis, G., Muroi, M., Richters, A., Winzker, M., Engbring, T., Tomassi, S., Watanabe, N., et al. (2017). Phenotypic Identification of a Novel Autophagy Inhibitor Chemotype Targeting Lipid Kinase VPS34. Angew. Chemie - Int. Ed. 56, 8153–8157.

Ronan, B., Flamand, O., Vescovi, L., Dureuil, C., Durand, L., Fassy, F., Bachelot, M.-F., Lamberton, A., Mathieu, M., Bertrand, T., et al. (2014). A highly potent and selective Vps34 inhibitor alters vesicle trafficking and autophagy. Nat. Chem. Biol. 10, 1013–1019.

Shang, J., Wan, Y., Luo, C., Ye, G., Geng, Q., Auerbach, A., and Li, F. (2020). Cell entry mechanisms of SARS-CoV-2. Proc. Natl. Acad. Sci. U. S. A. 117.

Silvas, J.A., Jureka, A.S., Nicolini, A.M., Chvatal, S.A., and Basler, C.F. (2020). Inhibitors of VPS34 and lipid metabolism suppress SARS-CoV-2 replication. bioRxiv 30307.

Vijgen, L., Keyaerts, E., Moës, E., Maes, P., Duson, G., and Van Ranst, M. (2005). Development of one-step, real-time, quantitative reverse transcriptase PCR assays for absolute quantitation of human coronaviruses OC43 and 229E. J. Clin. Microbiol. 43, 5452–5456.

Wei, J., Alfajaro, M., Hanna, R., DeWeirdt, P., Strine, M., Lu-Culligan, W., Zhang, S., Graziano, V., Schmitz, C., Chen, J., et al. (2020). Genome-wide CRISPR screen reveals host genes that regulate SARS-CoV-2 infection. bioRxiv 2020.06.16.155101.

Ye, Z.W., Yuan, S.F., Yuen, K.S., Fung, S.Y., Chan, C.P., and Jin, D.Y. (2020). Zoonotic origins of human coronaviruses. Int. J. Biol. Sci. 16, 1686–1697.

Zhou, P., Yang, X. Lou, Wang, X.G., Hu, B., Zhang, L., Zhang, W., Si, H.R., Zhu, Y., Li, B., Huang, C.L., et al. (2020). A pneumonia outbreak associated with a new coronavirus of probable bat origin. Nature 579, 270–273.

Zhu, N., Wang, W., Liu, Z., Liang, C., Wang, W., Ye, F., Huang, B., Zhao, L., Wang, H., Zhou, W., et al. Morphogenesis and cytopathic effect of SARS-CoV-2 infection in human airway epithelial cells. Nat. Commun. 1–8.

Zhu, Y., Feng, F., Hu, G., Wang, Y., Yu, Y., Zhu, Y., and Xu, W. (2020). The S1 / S2 boundary of SARS-CoV-2 spike protein modulates cell entry pathways and transmission. bioRxiv.

